# Genetic background influences MAFA^S64F^-mediated diabetes penetrance in male mice

**DOI:** 10.1101/2025.05.20.653758

**Authors:** Zachary Loyd, Dongsoo Lee, Mallory Maurer, Lizabeth Buzzelli, Jin-Hua Liu, Garrett Reynolds, Mark P. Keller, Jean-Philippe Cartailler, Mark A. Magnuson, Roland Stein, Jeeyeon Cha

## Abstract

Pancreatic β-cells require the coordinated expression of transcription factors such as MAFA to dynamically secrete insulin to maintain euglycemia. A naturally occurring mutation in MAFA (MAFA^S64F^) produces a long-lived variant protein which predisposes carriers to dichotomous conditions of either maturity (adult)-onset diabetes of the young (MODY) or hypoglycemia in a sex-dependent manner. Here we show that genetic background modulates disease penetrance in MafA^S64F/+^ male mice. Specifically, MafA^S64F/+^ males backcrossed to a C57/Bl6J (“C57”) background prevented dysglycemia, while C57 MafA^S64F/+^ males bred one generation to an SJL/J background (“mixed”) manifested overt diabetes, impaired insulin secretion, and accelerated β-cell senescence. MafA protein levels, phosphorylation status, and target gene expression in C57 MafA^S64F/+^ male islets were more comparable to wildtype males. RNA sequencing of C57 MafA^S64F/+^ male islets revealed fewer differentially expressed genes than mixed background male islets, including transcriptional signatures of β-cell senescence. In addition, retinoic acid signaling, another signature of cellular aging, was uniquely downregulated in C57 MafA^S64F/+^ male islets. Indeed, core retinoic acid signaling receptor RARα and known target genes were downregulated in C57 MafA^S64F/+^ male islets. CUT&RUN mapping revealed that MafA directly impacted retinoic acid receptor alpha *Rara* expression. In sum, these data show that genetic factors can impact unique pathways to modulate disease penetrance in mice modeling MAFA-MODY.

**Article Highlights:**

- Genetic background in mice or ancestry in humans can profoundly influence diabetes susceptibility.
- We asked whether diabetes penetrance in our robust mouse model of MAFA-MODY on a mixed genetic background was impacted when backcrossed to C57/Bl6J (“C57”).
- Here we show that C57 MafA^S64F/+^ males have wildtype glucose-sensing properties, compared to overt diabetes which could be reproducibly re-created on a mixed background. Further analysis disclosed more muted gene expression changes in C57 MafA^S64F/+^ male islets including unique downregulation of senescence and retinoic acid signaling.
- These results illustrate the influence of genetic landscape on β-cell function in response to a pathogenic MODY variant.

## INTRODUCTION

MafA, the large V-Maf avian <u>m</u>usculo<u>a</u>poneurotic fibrosarcoma transcription factor (TF), is a pancreatic islet β-cell-enriched protein that is essential for activating transcriptional programs promoting maturation (*1, 2*). Compromised levels of mouse MafA (*3–8*) or human MAFA (*9, 10*) also diminishes β-cell function. For example, genetic knockout models of pancreatic *MafA* manifest impaired glucose tolerance in male and female mice (*8*), while human *MAFA* expression is reduced in islets recovered from male and female donors with diabetes (*9*). Conversely, inducing *MafA* expression in MafA^Low^, non-glucose responsive neonatal rat islets increases glucose-stimulated insulin secretion (GSIS) (*11*), while acute introduction of human *MAFA* alone in mouse and human stem cells directly promotes insulin production and β-cell maturity markers (*11–13*). Collectively, these results demonstrate a fundamental role for MafA in β-cell function and identity.

Pathogenic mutations in pancreatic islet-enriched TFs including MAFA can cause heritable, monogenic forms of islet dysfunction in adults (*14*). A *MAFA* missense mutation (p.Ser64Phe, c.191C>T) in male carriers develop sex-dependent diabetes, while female carriers develop persistent hyperinsulinemic hypoglycemia due to the production of non-syndromic, insulin-secreting neuroendocrine tumors (insulinomatosis) (*15*). The resultant MAFA^Ser64Phe^ (termed MAFA^S64F^) variant protein impairs protein phosphorylation, which impacts both transcriptional activity and protein stability (i.e., t_1/2_ of MAFA is normally only ∼30 minutes, compared to many hours in MAFA^S64F^) (*15*). A mouse model harboring this clinically relevant mutation in the endogenous *MafA* gene (termed MafA^S64F/+^) on a mixed genetic background (C57/Bl6J and SJL/J) mimicked aspects of the heterogeneous, sex-dependent phenotypes: female MafA^S64F/+^ mice were persistently hypoglycemic, while male MafA^S64F/+^ mice showed impaired glucose tolerance due to widespread, premature β-cell aging and senescence (*16*). Introduction of this protein in a male human β-cell line also drove accelerated cellular aging (*16*), implying that the regulatory mechanisms are conserved between mice and humans.

Genetic background in mice (*17–20*) or ancestry in humans (*21–23*) can profoundly influence diabetes susceptibility. Common genetic variants have been identified to modify disease risk and clinical presentation in monogenic diseases including diabetes (i.e., MODY) (*24–27*). These observations prompted our current study to determine if MafA^S64F/+^ diabetes penetrance was impacted when backcrossed to the C57/Bl6J (“C57”) background. C57 was chosen since these mice are susceptible to glucose intolerance under conditions of stress (*17–20*). Here we show that C57 MafA^S64F/+^ males have essentially wildtype (WT) glucose-sensing properties by 6 weeks of age, compared to overt diabetes which could be re-created on a mixed background. Further analysis of C57 and mixed MafA^S64F/+^ male islets disclosed more muted gene expression changes in the C57 background as well as downregulation of β-cell senescence and retinoic acid signaling pathways associated with aging. These results illustrate the influence of genetic landscape on β-cell function in response to a pathogenic MODY variant.

## METHODS

### Mice

MafA^S64F/+^ heterozygous mice were originally generated using CRISPR/Cas9 targeting by the University of Michigan Transgenic Core on a mixed SJL/J and C57/Bl6J mouse strain (*16*). Mice were backcrossed eight generations with C57/Bl6J mouse strain (Jackson Lab) to generate C57 MafA^S64F/+^ mice (Transnetyx Genetic Monitoring, >95% C57Bl/6J). These C57 MafA^S64F/+^ mice were bred one time with SJL/J mice (Taconic) to produce the mixed genetic background strain (F1 generation, roughly 50:50 contribution of SJL/J and C57). Mice of both sexes were used in this study. All animal studies were reviewed and approved by the Vanderbilt University Institutional Animal Care and Use Committee. Mice were housed and cared for according to the Vanderbilt University Department of Animal Care and the Institutional Animal Care and Use Committee of Animal Welfare Assurance Standards and Guidelines.

### Intraperitoneal glucose tolerance testing and serum hormone measurements

Glucose tolerance testing was performed on male and female MafA^WT^ and MafA^S64F/+^ littermate mice (n = 6-16) after afternoon intraperitoneal injection of D-glucose (2 mg/g body weight) prepared in sterile PBS (20% w/v) preceded by a 5-hour fast (i.e. 8AM to 1PM). Afternoon insulin tolerance tests were conducted by intraperitoneal injection of 0.5 IU/kg body weight insulin (Novolin, regular human insulin, recombinant DNA origin) into mice (n = 3-5) fasted for 6 hours. Blood glucose was measured using a FreeStyle glucometer (Abbott Diabetes Care) before (0 minutes) and at 15, 30, 60, and 90 minutes following injection. Serum insulin and C-peptide was measured by Ultrasensitive Insulin ELISA (Mercodia) and C-peptide ELISA (Alpco), respectively.

### Mouse islet isolation, RNA isolation, and gene expression analysis

Mouse islets were isolated using collagenase P (Sigma) injected into the pancreatic duct, followed by a Histopaque 1077 (Sigma-Aldrich) gradient. Handpicked islets were isolated in standard RPMI-1640 medium (Thermo Fisher Scientific) supplemented with 10% FBS, L-glutamine, and penicillin-streptomycin, then stored frozen. Islet RNA was isolated using the RNAqueous Micro Total RNA Isolation Kit (Invitrogen). cDNA was generated using Superscript III reverse transcriptase (Invitrogen) by the oligo(dT) priming method (*16*). Real-time PCR assays were performed using the LightCycler FastStart DNA Master PLUS SYBR Green kit (Roche) and a LightCycler PCR instrument (Roche). The real-time PCR primers are listed in **Supplemental Table 1**. Gapdh or β-Actin was used to normalize the real-time PCR data which were analyzed using the ΔΔCt method (*8*).

### Cell culture and RNA isolation

Monolayer cultures of MIN6 mouse β-cells and HeLa cells were grown under conditions described previously (*4*). Transfection with scrambled control, MAFA^WT^ or MAFA^S64F^ constructs of confirmed sequences (*28*) was performed using Lipofectamine 2000 (Invitrogen) (average transfection efficiency 80%) before protein or RNA isolation. RNA was isolated 4 days posttransfection using the Trizol reagent (Fisher Scientific), while protein was isolated in RIPA buffer (*8*). cDNA synthesis and real-time PCR assays were performed as above (*16*).

### Western blotting and gel shift assay

MAFA protein levels were normalized to endogenous β-actin by immunoblotting with antibodies against MAFA (Cell Signaling Technology, 79737) or β-actin (MilliporeSigma, MAB1501). Horseradish peroxidase–conjugated anti-rabbit (31460, Thermo Fisher Scientific) or anti-goat (31402, Thermo Fisher Scientific) secondary antibodies were used at 1:5000 dilution. Immunoblots were quantified with ImageJ (NIH). Gel shift assays were performed using 10 μg of nuclear protein extract and 200 fmol of biotin-labeled double-stranded human *INS* MAFA binding site probe alone or mixed with WT unlabeled competitor DNA, mutant unlabeled competitor DNA, or MAFA antibody (Bethyl Laboratories, Product # A700–067) in a total of 20 μL reactions (Lightshift Chemiluminescent EMSA Kit, Thermo Fisher Scientific) containing 1X binding buffer, 2.5% glycerol, 5 mM MgCl, 50 ng/μL poly(dI-dC) (Thermo Fisher Scientific), and 0.05% NP-40 (Sigma-Aldrich). These reactions were electrophoresed through a 6% precast DNA retardation gel in 0.5% Tris-borate-EDTA buffer (TBE, Thermo Fisher Scientific) at 100 V for 1.5 h.

### Immunohistochemical analyses

Mouse pancreata were fixed overnight at 4°C in 4% paraformaldehyde (Electron Microscopy Services) in PBS, and pancreata were embedded in either Tissue-Plus OCT (Thermo Fisher Scientific) or paraffin wax. Sections of rodent pancreata were cut at 6μm thickness. The paraffin sections were deparaffinized and rehydrated before citrate buffer–based antigen retrieval. Sections were then made permeable by 0.5% Triton in PBS treatment for 10 minutes. Following blocking with 0.5% BSA in PBS for 120 minutes, the primary antibodies were applied overnight at 4°C. Primary antibodies are listed in **Supplemental Table 1**. Species-matched antibodies conjugated with the Cy2, Cy3, or Cy5 fluorophores were used for secondary detection (1:1000; Jackson ImmunoResearch). DAPI was used for nuclear staining (Southern Biotech). Immunofluorescence images were obtained using the Zeiss Axio Imager M2 widefield microscope with ApoTome.

### Senescence-associated β-galactosidase (SA-β-gal) staining

Pancreata were snap frozen in O.C.T. and cryosections were prepared at 16μm thickness. SA-β-gal activity staining was performed at pH 6.0 (*16*) (Cell Signaling). To compare the intensity of SA-β-gal staining, sections from different genotypes were processed on the same slide. Staining reactions were developed for 18 hours at 37°C, then quenched by 3x PBS washes (pH 7.4). These slides were subject to immunostaining for insulin by fixing in 4% paraformaldehyde for 45 minutes, permeabilized with Tris-buffered saline with 0.2% Triton in PBS for 15 minutes, blocked in 2% normal donkey serum/1% BSA in PBS and incubated overnight with guinea pig anti-insulin (1:500, Fitzgerald 20-IP35) at 4°C. HRP-conjugated secondary antibodies were incubated on slides for 1 hour and detected with the DAB+ chromogen kit (DAKO). After washing, slides were mounted and imaged by brightfield microscopy.

### RNAscope and quantification

The RNAscope® Multiplex Fluorescent Reagent Kit V2 (ACD) was used following manufacturer’s instructions and modified for mouse pancreata as previously described (*29*). RNAscope images from 6 week old mice were generated using a Leica DMi8 inverted microscope at 40X objective and analyzed using QuPath (*30*). Regions of interest (ROI) around islets were defined based on insulin-positive staining. Stardist (*31*), a deep-learning-based nuclei detection model, was used to detect nuclei and outline the cell. Max entropy or triangle auto-threshold methods and image down-sampling (factor of 2) was applied to determine the threshold for each channel with an RNA probe. RNA speckles were detected using the subcellular detection feature in QuPath with the calculated thresholds. Percentage of β-cells positive for *Cdkn1a* was calculated in QuPath and exported to Prism for statistical analysis.

### Bulk RNA-sequencing analysis of mouse islet cells

Bulk RNA-sequencing was performed on mouse islets isolated at 6 weeks of age. RNAqueous Micro Total RNA Isolation Kit (Invitrogen) was used to isolate total RNA, and RNA quality was analyzed on an Agilent 2100 Bioanalyzer. Samples with RIN >8 were used for sequencing. The cDNA libraries were constructed, and paired-end sequencing was performed on an Illumina NovaSeq6000 (150-nucleotide reads). The generated FASTQ files were processed and interpreted using the Genialis visual informatics platform (https://www.genialis.com) (*16*). DESeq2 was used for differential gene expression analyses and statistical comparison (*32*). Poorly expressed genes (i.e., expression count summed over all samples below 10) were filtered out from the differential expression analysis input matrix. Sequencing data is available in the National Center for Biotechnology Information (NCBI) Gene Expression Omnibus (GEO) database (accession no. GSE299343).

### Position Weight Matrix (PWM) Motif Search Protocol

Putative MAFA responsive elements (MAREs) were identified in the *Rara* gene promoter and other upstream regulatory regions by scanning genomic sequences for Position Weight Matrix (PWM) motifs. The MAFA PWM was obtained from the JASPAR database (matrix ID MA1521.2), representing the 13 bp core MARE consensus derived from ChIP-seq binding data (*33*). The position frequency matrix was converted to a log-odds PWM with pseudocounts of 0.8 using TFBSTools (*34*). Genomic sequences were scanned on both strands using TFBSTools::searchSeq(), and matches scoring at or above 80% of the maximum possible PWM score were retained as candidate binding sites. The transcription start site (TSS) was defined as the 5’-most position of a representative transcript from the UCSC Gene annotation (mm39 assembly); one TSS per gene symbol was used. Three non-overlapping search regions were defined relative to each TSS: proximal promoter (TSS -200bp to +1000bp), distal promoter (TSS +1kb to +5kb), and enhancer (TSS +5kb to +100kb). PWM relative scores were tiered as follows: 80-85% (weak/marginal candidate), 85-90% (moderately strong candidate), and 90%+ (strong candidate with high confidence match to consensus sequence).

### CUT&RUN assay and analysis

CUT&RUN was performed on 500,000 dispersed mouse MIN6 cells per condition using the CUTANA ChIC/CUT&RUN protocol v3.1 (Epicypher). Nuclei were extracted with nuclear extraction buffer (20 mM HEPES-KOH [pH 7.9]; 10 mM KCl; 0.1% Triton X-100; 20% glycerol; 1 mM MnCl2; 0.5 mM spermidine; 1uM protease inhibitor; Thermo Fisher Scientific) for 10 minutes on ice and immobilized onto Concanavalin-A beads (EpiCypher). After blocking and washes, samples were incubated with 0.5 μg of rabbit anti-MAFA (Cell Signaling 79737) or rabbit anti-IgG (EpiCypher 13-0042) antibodies overnight at 4°C. pAG-MNase (EpiCypher) was added to nuclei (1:20) and incubated at room temperature for 10 minutes. Targeted chromatin digestion was induced by adding 100 mM CaCl_2_ and nutating for 2 hours at 4°C. DNA fragments were purified using the CUTANA ChIC/CUT&RUN kit. DNA was resuspended in 0.1M∼ Tris-EDTA buffer solution and used for library preparation with the CUTANA CUT&RUN Library Prep Kit (EpiCypher, 14-1001), according to the version 1 (v1) manual. Libraries were sequenced as PE150 reads on the NovaSeq platform. All libraries had > 15 million reads and were processed using the nf-core/cutandrun workflow, v3.1 (*35*). The pipeline performs adapter trimming, alignment to the GRCm38 reference genome, filtering against the mm10 blacklist regions, spike-in normalization, and peak calling using MACS2. The workflow also generates IGV sessions and all track data, which allows for interactive visualization and exploration. Quality was confirmed by fragment length distribution and fingerprint plots. Nearest gene analysis was performed with peakScout (*36*). Coverage plots were created using ggcoverage (*37*). Sequencing data were deposited in the National Center for Biotechnology Information (NCBI) Gene Expression Omnibus (GEO) database (accession no. GSE298664).

### Statistical Analysis

Data are expressed as the mean ± SEM. Statistical analysis was performed using GraphPad Prism 9.5.0 (GraphPad Software Inc.). The differences between groups were analyzed by unpaired 2-tailed Student’s t test or 2-way ANOVA, as indicated. Differences were considered to be statistically significant at P < 0.05.

## RESULTS

### Glucose tolerance is largely unaffected in C57 MafA^S64F/+^ males at 6 weeks of age, while those on a mixed background manifest overt diabetes and impaired insulin secretion

As seen clinically, MafA^S64F/+^ mice on a mixed genetic background (C57/Bl6J and SJL/J) were previously found to have distinct metabolic phenotypes (*16*). Here we find that impaired glucose tolerance of MafA^S64F/+^ males was essentially eliminated by 6 weeks of age when backcrossed eight generations to the C57/Bl6J (C57) genetic background (**Fig. 1A**, C57). In contrast, the F1 generation of C57 MafA^S64F/+^ males bred once with parental SJL/J mice were both glucose intolerant and hyperglycemic (**Fig. 1B**, mixed) due to impaired insulin and C-peptide secretion (**Fig. 1C-D**). The newly generated F1 mixed background mice were glucose intolerant, like the earlier MafA^S64F/+^ males (**Fig. 1B**) (*16*) and showed transient hypoglycemia preceding impaired glucose tolerance at 5 weeks of age even on an extended wean protocol (**Supp Fig. 1A-B**). Moreover, this mixed MafA^S64F/+^ mouse line had comparable weight gain as well as intact peripheral insulin tolerance in relation to littermate wild type (WT) mice (**Supp Fig. 1C-E**), all of which were properties of the previously reported mixed line (*16*). In contrast, MafA^S64F/+^ females on both mixed and C57 backgrounds were hypoglycemic as reported previously (*16*). Consequently, we principally focused our efforts on determining how the C57 background prevented glucose intolerance in MafA^S64F/+^ males.

**Figure 1.**
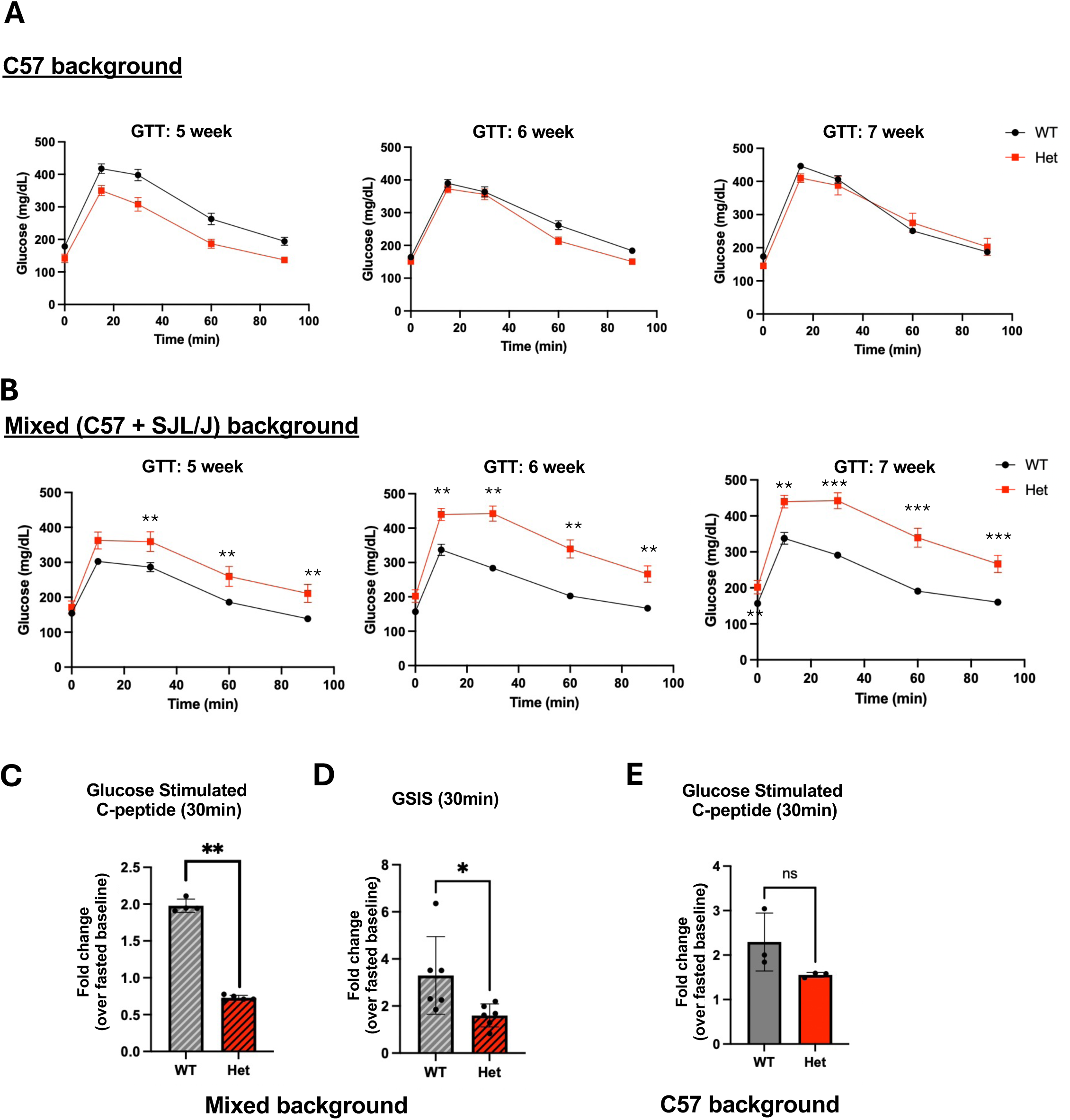
Glucose tolerance is largely unaffected in C57 MafA^S64F/+^ males, while those on a mixed background manifest frank diabetes and impaired insulin secretion. A. Fasted male animals underwent intraperitoneal glucose tolerance tests (GTT) at time points indicated. Male MafA^S64F/+^ heterozygous (Het) mice on a C57 background had no changes to glucose tolerance through adulthood compared to MafA^+/+^ (WT) male littermates. B. Het males on a mixed background had impaired glucose tolerance by 5 weeks of age which then progressed to overt diabetes by 7 weeks of age. **p < 0.01; ***p < 0.001. C-D. Stimulated in vivo C-peptide (C) and insulin (D) measurements showed reduction in Het males at 6 weeks of age on a mixed background. Two-tailed Student t test; *p < 0.05; **p < 0.01. E. Stimulated in vivo C-peptide measurements showed no significant changes between WT and Het males at 6 weeks of age on a C57 background.

### MafA^WT^ protein status and target gene expression is augmented in C57 MafA^S64F/+^ male islets compared to mixed background males

MafA function is dependent upon posttranslational phosphorylation events which couple two distinct regulatory processes: increased transactivation activity *and* ubiquitin-mediated degradation (*38*). *In vitro* analysis of MafA^S64F^ found that impaired phosphorylation at neighboring Ser65 (S65) prevented the ability of glycogen synthase kinase 3 (GSK3) to phosphorylate S61, T57, T53, and S49, which results in a protein with faster SDS-PAGE mobility, greater stability, and altered transactivation capacity (*38*). Here we show that the MafA^WT^ protein was visible in C57, but not mixed, background MafA^S64F/+^ male islets (**Fig. 2A**). Interestingly, the quantity of MafA^S64F^ protein did not appear to differ by genetic background (**Fig. 2B**). The expression of MafA and known MafA target genes were either less impacted (i.e., *Ins1*, *Ins2*) or unaffected (i.e., *Gck, Slc2a3*, *Slc30a8, MafA*) in C57 MafA^S64F/+^ male islets (**Fig. 2C**), presumably reflecting MafA^WT^ protein activity in this background. The relatively low level of total islet MafA (WT+S64F) protein in relation to WT mice reflects the ability of MafA^S64F^ to reduce endogenous mRNA production (**Fig. 2C**) (*16*). Along these lines, low MafA protein levels were also evident in human MAFA^S64F^ carriers (*15*).

**Figure 2.**
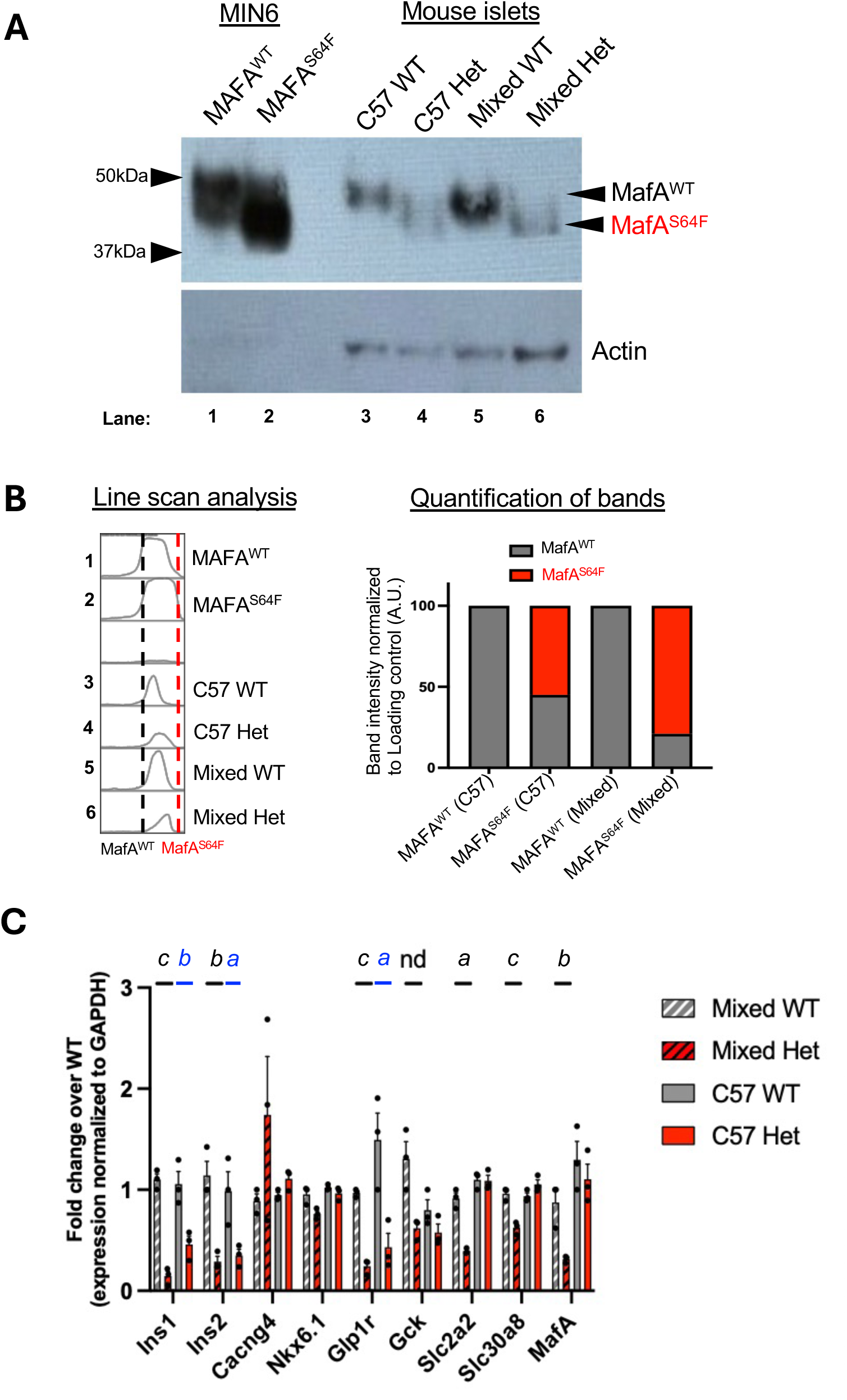
MafA protein status and activity are augmented in islets of C57 MafA^S64F/+^ male islets compared to those from a mixed background. A. Left lanes (1-2), representative western blotting on MIN6 nuclear extract transfected to express either MafA^WT^ or MafA^S64F^ shows faster migration in mutant MAFA due to impaired posttranslational modification by phosphorylation. Right lanes (3-6), Isolated mouse islets from each genotype and background (150 islets per lane) showed detectable levels of phosphorylated MafA^WT^ in C57 background, but relative uniformity of MafA species with faster migration and impaired phosphorylation (MafA^S64F^) in the mixed background. B. Quantification of western blotting bands by line scan analysis shows greater proportion of phosphorylated MafA^WT^ species (gray) in Het male islets from the C57 background compared to the mixed background. C. qPCR of known MafA target genes in male islets between backgrounds. Blue letters, comparison between C57 WT and C57 Het islets; Black letters, comparison between mixed WT and mixed Het. ***a***, p< 0.05; ***b,*** p < 0.01; ***c,*** p < 0.001; **nd,** no difference. Mean ± SEM. n=3-4 mice per group.

### Unbiased RNA sequencing illustrates how differently MafA^S64F^ regulates gene expression in C57 and mixed MafA^S64F/+^ male islets

Bulk RNA sequencing was performed on MafA^WT^ and MafA^S64F/+^ male islets at 6 weeks of age to assess dominant gene expression patterns between these groups. Notably, the 6 week time point in MafA^S64F/+^ males represents when distinct differences in male phenotypes between backgrounds is observed (**Fig. 1A-B**). Male C57 MafA^S64F/+^ islets had 2,097 differentially expressed genes (DEGs, 616 up and 1481 down) compared to male WT islets, while a greater number of DEGs were found on the mixed background MafA^S64F/+^ mice (3,366: 1505 up and 1861 down) (**Fig. 3A**). Of these, only 386 upregulated and 621 downregulated DEGs were similarly impacted between the genetic backgrounds. Unbiased clustering identified that the molecular signatures between MafA^WT^ and MafA^S64F/+^ males on a C57 background were more similar than those on a mixed background (**Fig. 3B**). In addition, GO pathway analysis showed that the up- and down-regulated DEGs separated into distinct GO pathways in C57 and mixed MafA^S64F/+^ male islets (**Fig. 3C**).

**Figure 3.**
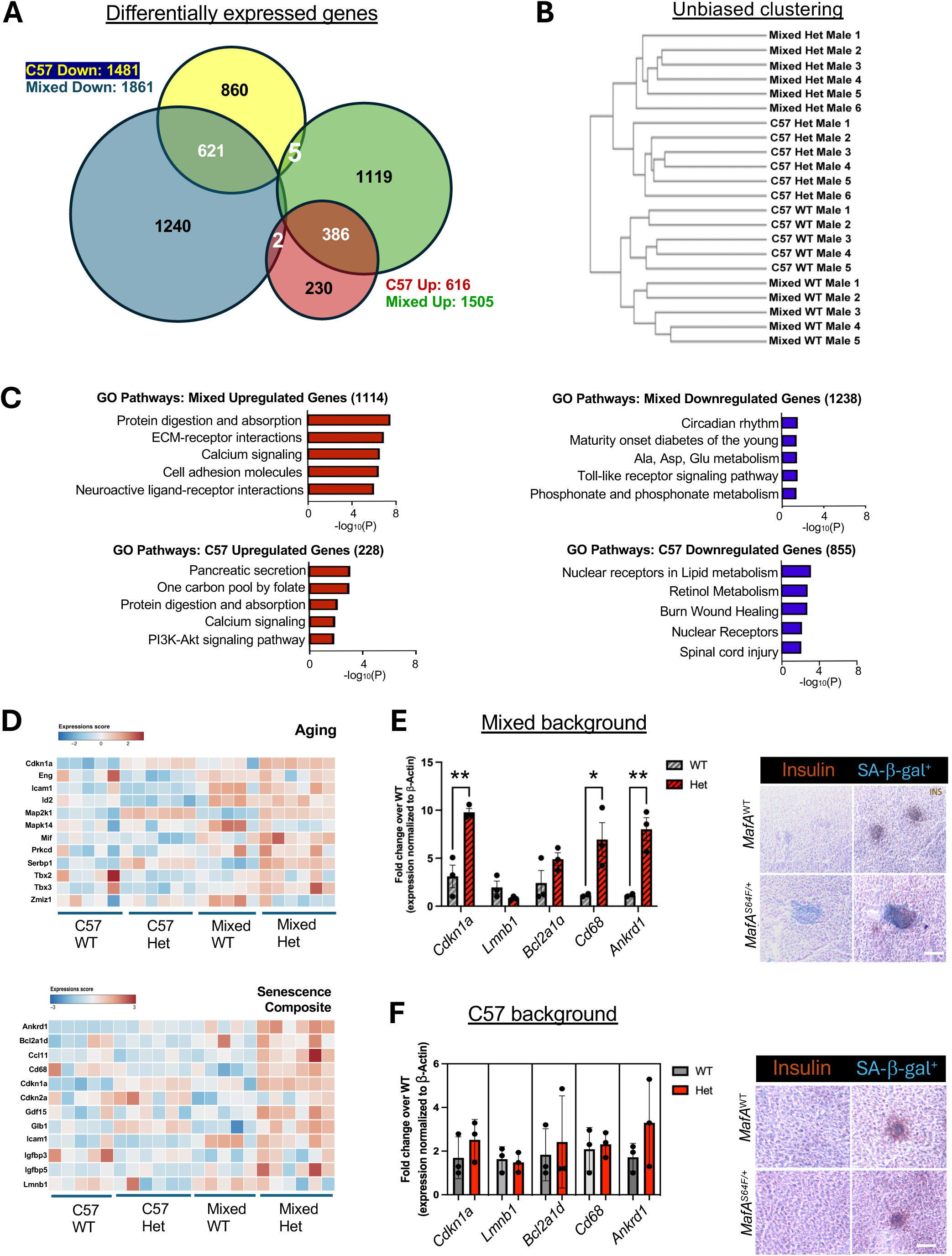
Male C57 MafA^S64F/+^ islet gene expression is more similar to MafA^WT^ islets. A. Venn diagram illustrates the total number of RNA-seq-identified DEGs up- or down-regulated between 6-week-old WT and Het male islets of each background. B. Unbiased hierarchical clustering showed congruent organization within genotypes and genetic background. C. Gene Ontology (GO) molecular function analysis (p < 0.05) of the genes uniquely upregulated in each genetic background in male Het islets. D. Heatmaps showing aging/senescence enrichment in mixed, but not C57, Het male islets. Each column represents islets from a unique animal. E. Left, qPCR of senescence signature genes are enriched in Het islets from a mixed background. Right, senescence associated beta gal staining (SA-β-gal) is enriched in male Het islets from the mixed background. *p < 0.05, **p < 0.01. Scale bar, 20μm. F. Left, qPCR of senescence signature genes are not enriched. Right, SA-β-gal staining not detectable in male Het islets from the C57 background at 6 weeks of age.

### Senescence gene signatures are abrogated in male C57 MafA^S64F/+^ islets

GO analysis and STRING protein association network analysis (*39*) of the DEGs of MafA^S64F/+^ male islets defined a distinct set of genetic background-specific pathways (**Fig. 3C**). As expected, cellular aging pathways were found in mixed MafA^S64F/+^ male islets, which represent earlier reported signatures (*16*). However, these pathways were attenuated in the C57 MafA^S64F/+^ males (e.g., see pancreatic secretion, protein digestion and absorption, ECM receptor interactions, cell adhesion, and calcium signaling pathways). In line with this, senescence gene signatures enriched in the mixed background MafA^S64F/+^ male islets (*16*) were largely muted in the male C57 MafA^S64F/+^ islets (**Fig. 3D-F**). As previously reported (*16*), islet senescence signatures were not seen in female MafA^S64F/+^ islets (**Supp Fig. 2**). Furthermore, hallmark molecular changes associated with cellular senescence such as enrichment of senescence associated-β-galactosidase (SA-β-gal) staining, DNA damage response such as 53BP1, and impaired nuclear integrity such as by downregulation of LaminB1, found in mixed background MafA^S64F/+^ males, were not detected or unchanged in C57 MafA^S64F/+^ male islets (**Figs. 3E-F, 4A-B**). Collectively, these results revealed that glucose intolerance (**Fig. 1**) and accelerated islet aging was prevented in C57 MafA^S64F/+^ male islets.

**Figure 4.**
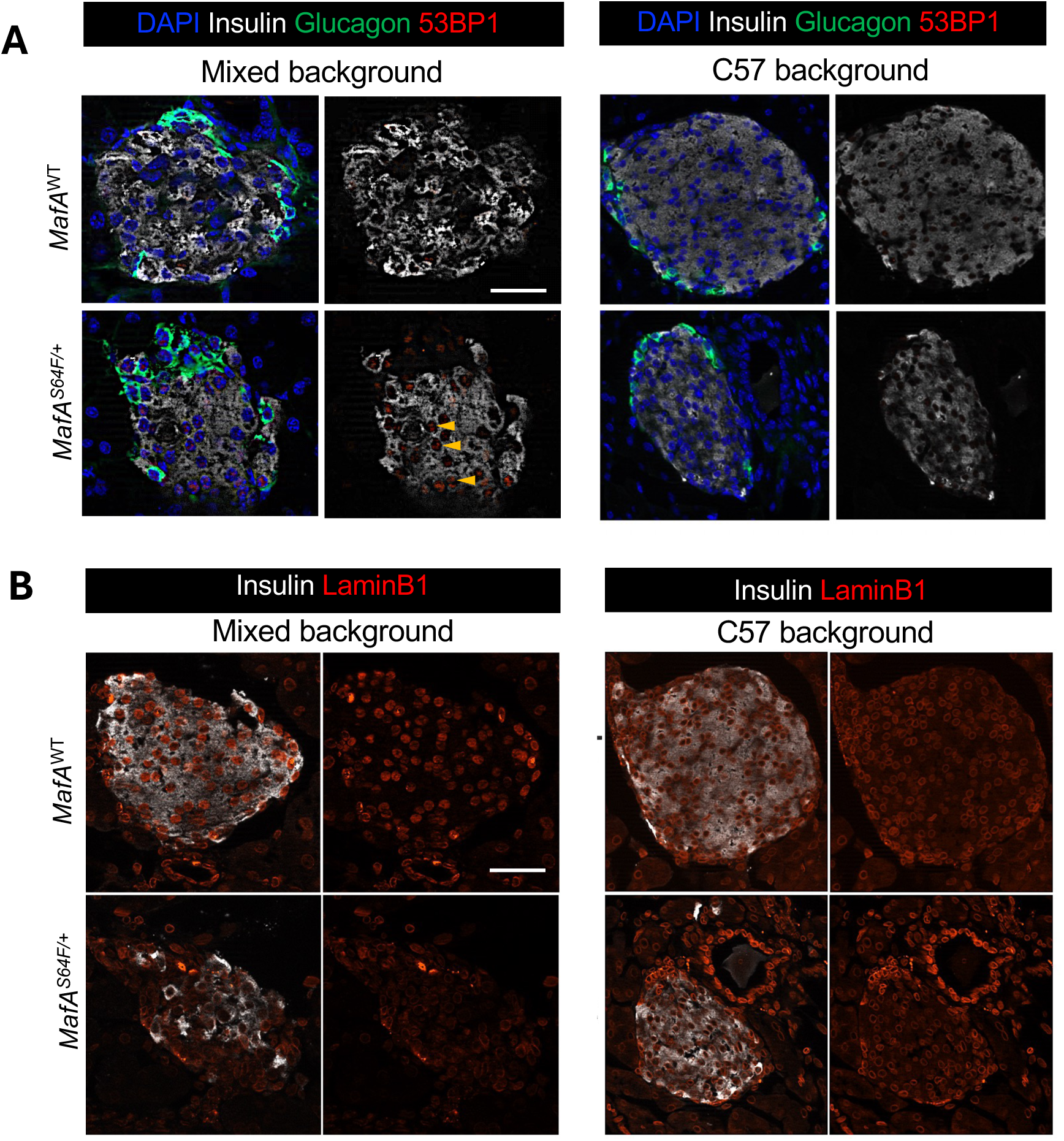
Islet senescence is abrogated in MafA^S64F/+^ male islets on a C57 background. A-B. Immunostaining of senescence markers show enrichment of 53BP1 (A) and relative loss of nuclear LaminB1 (B) in the mixed, but not C57, background. Scale bar, 50μm. Arrowhead, positive stain for 53BP1.

### C57 MafA^S64F/+^ male islets show unique downregulation of core retinoic acid signaling components

The activity of mixed male MafA^S64F/+^ islets is presumably mediated by unique GO pathways associated with cellular aging and MODY enriched in this sex and background (**Fig. 3C**). Circadian rhythm was another downregulated pathway in the mixed background males. Indeed, oscillatory insulin secretion is synchronized with nutritional intake (*40–43*), thus the circadian dysregulation in β-cells observed in this context could be additive to accelerated aging to impair dynamic insulin secretion and promote dysglycemia (*42–44*). In addition, downregulation of nuclear receptors and retinol metabolism genes was observed by GO analysis in C57 MafA^S64F/+^ islets (**Fig. 3C**). In line with this, persistent retinoic acid signaling through retinoic receptors RAR or RXR drives cell cycle arrest and senescence (*45–47*), and exogenous retinoic acid induces senescence programming via upregulation of p21 and p16 cell cycle inhibitors (*48–50*).

Importantly, the core retinoic acid signaling components *Retinoic acid receptor (Rar) a, Rarb*, and *Retinol binding protein (Rbp) 7* were only downregulated in C57 MafA^S64F/+^ male islets (**Fig. 5A**). MafA CUT&RUN analysis was performed in mouse Min6 β-cells to determine whether retinoic acid signaling components could be gene targets of endogenous MafA. We expect that this will also reveal MafA^S64F^ binding in β-cells, as this mutation resides in the transactivation, and not DNA binding, domain. Gel shift assay showed that MafA^WT^ and MafA^S64F^ have similar *cis*-element binding ability when transfected in HeLa cell nuclear extracts (**Supp Fig. 3**) (*8*).

**Figure 5.**
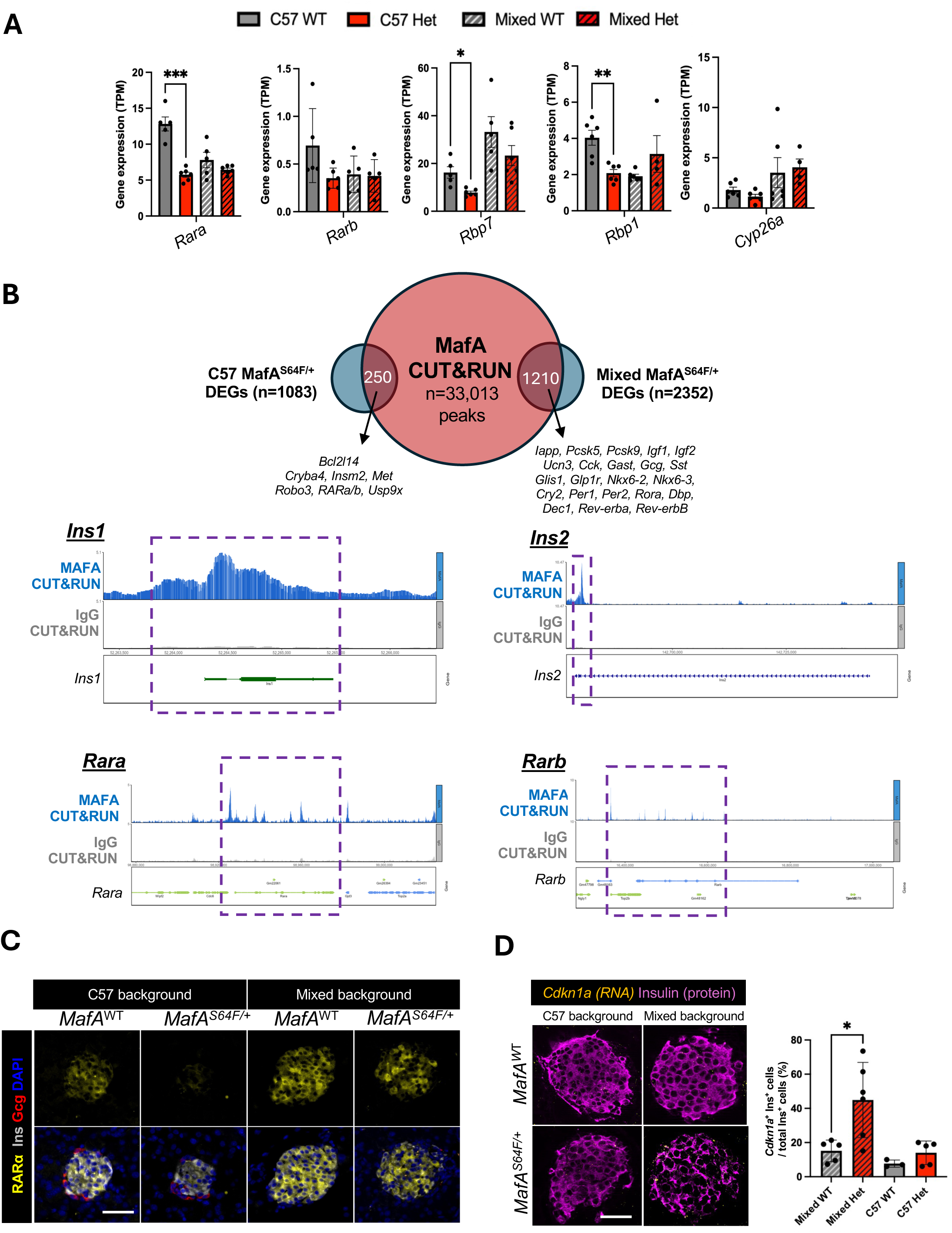
MafA directly modulates downregulation of retinol signaling in C57 MafA^S64F/+^ male islets. A. Expression of retinoic acid receptors signaling components and known target genes Rbp1 and Cyp26a are altered specifically in C57 MafA^S64F/+^ (Het) male islets. TPM±SEM. *p < 0.05,**p < 0.01. ***p < 0.001. n=5-6 animals/group. B. Top, DEGs identified uniquely in Male Het islets from a mixed background (1238 + 1114 = 2352 genes) versus those uniquely from a C57 background (228 + 855 = 1083 genes) were then overlayed with peaks identified by endogenous MafA CUT&RUN in mouse MIN6 cells (n=33,013 peaks). Of these, 250 DEGs uniquely enriched in a C57 background overlapped with a MafA CUT&RUN peak, while 1210 DEGs were uniquely enriched in a mixed background overlapped with a MafA CUT&RUN peak. Bottom, UCSC Genome Browser tracks showing genomic regions associated with endogenous MafA CUT&RUN peaks near known targets *Ins1* and *Ins2* (positive controls), and candidate genes *Rara* and *Rarb*. MafA CUT&RUN enriched peaks are highlighted in dashed boxes, and regulated genes are depicted below IgG control tracks. C. Immunostaining of RARα show downregulation in the C57 but not mixed background MafA^S64F/+^ males. Scale bar, 20μm. D. Top, RNAScope for *Cdkn1a* encoding p21 shows enrichment in Het islets on a mixed background in β-cells. Bottom, Quantification of *Cdkn1a*^+^Ins^+^ cells. *p < 0.05.

Gene-annotated MafA CUT&RUN peaks (n=33,013 peaks) were overlaid with the DEGs uniquely identified in MafA^S64F/+^ males of each genetic background (**Fig. 5B**). Mixed background MafA^S64F/+^ male islets yielded 1210 putative targets (**Fig. 5B**). Importantly, these included genes associated with β-cell identity and islet hormones such as urocortin 3 (Ucn3), glucagon (Gcg), somatostatin (Sst), cholecystokinin (Cck), insulin (Ins), and gastrin (Gast); islet-enriched transcription factors and receptors such as Glis1, Glp1r, Nkx6.2, Nkx6.3, and Smarca1; and circadian regulators Period1/2 (Per1/2), Crytochrome2 (Cry2), RAR-related orphan receptor α (Rora), D-site albumin promotor binding protein (Dbp), basic helix-loop-helix family member (Bhlhe40/Dec1), and nuclear receptor subfamily 1 group D member 1 (Rev-erba and Rev-erbb), which sets up multiple highly effective feedback loops that promotes dynamic insulin secretion (*18, 37, 38, 51, 56–59*). Collectively, these results suggest several avenues of dysregulation in mixed background MafA^S64F/+^ male islets.

C57 MafA^S64F/+^ male islets yielded 250 unique putative targets of MafA (**Fig. 5B**). Notably, these targets included factors in the senescence-associated prosurvival Bcl2 family (ie., Bcl2l14) and retinoic acid receptor (RARα and RARβ) signaling. Consistent with these findings, *Rara* houses a moderately strong candidate MARE (Maf responsive binding element) in its distal (+1000 to +5000bp) promoter region. Immunostaining also revealed downregulation of RARα protein in C57 MafA^S64F/+^ male islets (**Fig. 5C**), and gene expression studies showed downward trends of classical RARα signaling targets *Rbp1* and *Cyp26a1* (**Fig. 5A**). Furthermore, a significant reduction in RARα target *Cdkn1a* (encoding p21) was seen in C57 MafA^S64F/+^ islet β-cells compared to mixed background MafA^S64F/+^ β-cells (**Fig. 5D**). Thus, we conclude that MafA can directly regulate *Rara* expression in C57 MafA^S64F/+^ β-cells to attenuate *Cdkn1a* induction and β-cell senescence in MafA^S64F/+^ islets.

## DISCUSSION

Genetic ancestry is widely accepted to influence disease penetrance (*21–23*). Even in purported monogenic disease, there is increasing recognition that genetic factors can play modulating roles beyond the causative mutation (*25–27*), and a recent study has shown that common genetic variants can influence monogenic HNF-MODY diabetes risk to range from 11% to 81% (*24*). Genetic background influences can more easily be mechanistically addressed in monogenic diabetes than in polygenic type 1 (T1D) and type 2 (T2D) diabetes. As a result, here we investigated the influence that genetic background could have on the MAFA-MODY variant and found a profound impact on the male diabetic phenotype in mice.

Patient carriers of the MAFA^S64F^ missense mutation (*15*) manifest sex-dependent glycemic disorders: men show adult-onset diabetes while women primarily manifest hypoglycemia with non-syndromic insulinomatosis (*15*). Since this original report, a second human MAFA missense mutation in the transactivation domain (i.e., MAFA^T57R^) has been identified to block the actions of GSK3 and, as expected, promotes MAFA protein stability and cosegregates with insulinomatosis and MODY-like diabetes (*51*). A mouse model harboring the S64F mutation in the endogenous *Mafa* gene, which prevented phosphorylation of the priming kinase at MAFA^S65^ and prevents GSK3 actions, recapitulated the sex-dependent responses seen in patients (*16*), providing mechanistic insights into the impact of this MafA variant *in vivo*, including a male sex-dependent acceleration of islet senescence that impaired β-cell function. In fact, accelerated cellular aging is a common feature of islets from T1D and T2D donors, and is seen in mouse models of diabetes such as those with insulin resistance modeling T2D (*52*), NOD mice modeling T1D (*53*), and MafA^S64F/+^ males modeling MAFA-MODY (*16*).

Given increasing appreciation for genetic background effects in diabetes penetrance, efforts to backcross MafA^S64F/+^ mice on the C57 background were examined. Here we show that the C57 genetic background interacts with this MafA mutation to influence the penetrance of the male dysfunction by: 1) preventing glucose intolerance and β-cell senescence; 2) allowing increased MafA^WT^ protein production, 3) yielding gene expression pattern similar to MafA^WT^ islets; and 4) downregulating retinoic acid signaling, an important mediator of β-cell maturity.

Spatiotemporally regulated retinoic acid signaling is a requisite for cellular differentiation of many cell types including islet β-cells. Specifically, a necessary role of retinoic acid signaling has been found in β-cell differentiation and function in mouse and human islets (*54, 55*), and it is also a critical component for stem-cell derived maturation of β-like cells for cell-based therapies (*56, 57*). However, persistent retinoic acid signaling and exogenous retinoic acid can activate cell cycle arrest and senescence programs (*45, 46*) via core cell cycle inhibitors (e.g., p21) across cell types (*48–50*). In fact, excess signaling through endogenous pancreas retinoids are reported to contribute to glucose intolerance (*58*). Here we show that downregulation of this accessory maturity pathway in C57 MafA^S64F/+^ mice appears to dampen the expression of the common aging driver p21.

Conversely, in the mixed background, we uncovered an impact of MafA^S64F^ on islet circadian regulators, identifying a novel link between β-cell maturity driver MafA, cellular aging, and circadian entrainment. Interestingly, we find that male MafA^S64F/+^ islets on the mixed background show marked downregulation of not only direct circadian regulators (e.g. *Per1*, *Per2*, *Cry2)* but also amplification factors in complementary circadian circuits such as *Dbp, Rora, Bhlhe40/Dec1, Rev-erb* and *Nfil3* (*42–44*). Interestingly, the expression of core circadian activators Bmal and Clock was unchanged between genotypes and genetic background, affirming that the genetic-background related circadian dysregulation engaged regulators rather than the core Clock:Bmal1 components themselves. A recent study in human stem-cell derived β-cells deficient in *Bhlhe40/Dec1* showed impaired MafA periodicity, suggesting a novel circadian islet feedback loop (*59*), but confirmation of this relationship in *MafA^S64F/+^* islets requires further study. Notably, several senescence mediators such as *Cdkn1a*, *Bcl2a1d*, *Ankrd1*, or *Icam1* enriched in mixed background MafA^S64F^ male islets, were not identified as MafA targets by CUT&RUN, suggesting that their regulation is through the influence of MafA^S64F^ on intermediate pathways such as circadian regulators or retinoic acid signaling.

MafB is another islet-enriched Maf family member with distinct temporal and cell-type expression from MafA (*4*). In mouse islets, MafB is the dominant family member expressed developmentally before MafA is expressed postnatally, while human MAFB is postnatally expressed alongside MAFA in β-cells. The analogous mutation in human MAFB (MAFB^Ser70Ala^, or MAFB^S70A^) in the transactivation domain also impairs phosphorylation and increases protein stability (*28*). Notably, it is one of several mutations which drives the clinical syndrome Multicentric Carpotarsal Osteolysis characterized by bony deformities and renal failure (*60*). The clinical impact of MAFB^S70A^ reflect the broader expression pattern of MAFB compared to MAFA (*1*). While the effect of this mutation on islet cell function is unclear, MAFB is essential for the development of human embryonic stem cell derived β-like Insulin^+^ cells (*61*), thus future studies comparing MAFB^S70A^ to MAFA^S64F^ would provide insights into its pathogenicity to adult β-cells.

A crucial next step is to investigate MAFA^S64F^ or MAFB^S70A^ in genetically modified human islets. However, our results raise the possibility that studies in male human islets may not show consistent phenotypes due to donor genetic background. In such a case, further investigations on the male phenotype may require models controlling for genetic background, such as stem cell derived β-cells, or a return to mouse models such as ours to investigate genetic background influences. In mice, genome-wide differential single nucleotide polymorphism (SNP) analysis between inbred genetic backgrounds can be pursued, and prioritization of high impact variants in genes involved in retinoic acid signaling, cellular aging, and circadian dysregulation would be interesting. Furthermore, identification of SNPs in genes which can impact MafA turnover such as kinases, ubiquitination mediators or proteasome components would also be interesting given the relative integrity of MafA^WT^ protein in C57 MafA^S64F/+^ mice. An unbiased forward genetic screen can mechanistically delineate specific genetic loci which can contribute to the dramatic differences in MafA expression and activity in MafA^S64F/+^ males between genetic backgrounds.

Nevertheless, our work here demonstrates the combinatorial impact of genetic background and biologic sex on molecular and functional β-cell responses to produce heterogenous responses (i.e., cellular senescence driving dysfunction vs. hypoglycemia) to a clinically relevant, stable variant of MafA. Our data strongly suggest that additional genetic regulators can modulate cellular responses to a pathogenic mutation and ultimately will improve strategies for diabetes risk stratification and therapies to optimize β-cell function in people at risk of diabetes.

## Acknowledgements

This research was performed using resources and/or funding provided in parts by NIH grants to JC (K08 DK132507); RS (R01 DK090570); Alan Attie and MPK (R01DK101573, R01DK102948, and RC2DK125961); and the Vanderbilt Diabetes Research and Training Center (DK20593). JC was also supported by a Doris Duke Physician Scientist Fellowship, Burroughs Wellcome Fund Career Award for Medical Scientists, and generous start-up funding from Vanderbilt University Medical Center. MPK is also supported by the University of Wisconsin-Madison Department of Biochemistry and Office of the Vice Chancellor for Research and Graduate Education with funding from the Wisconsin Alumni Research Foundation. The authors thank former and current members of the Cha lab for feedback on this manuscript.

## Guarantor Data Access and Responsibility Statement

JC is the guarantor of this work, and as such, had full access to all data in the study and is responsible for the integrity of the data and accuracy of data analysis.

## Author Contributions

ZL, DL, MAM, RS, and JC designed the study.

ZL, DL, MM, LB, JL, GR, JPC, and JC performed experiments.

ZL, DL, MM, MPK, JPC, and JC performed bioinformatic analysis.

ZL, DL, MM, LB, JL, GR, MPK, JPC, MAM, RS, and JC analyzed data.

ZL, DL, JPC, RS and JC wrote the manuscript.

ZL and DL are co-first authors due to their contributions in most of the experimental studies.

## SUPPLEMENTAL FIGURE AND TABLE LEGENDS

**Supplemental Figure 1. Mixed background male and female MafA^S64F/+^ show hypoglycemia while nursing, extended wean does not impact hyperglycemia in male Hets, and weight gain and insulin tolerance are similar between the genetic backgrounds**

A. Random blood glucose monitoring at 2-3 weeks of age while nursing shows similar mild, transient hypoglycemia in male and female Het mice on a mixed background. *p < 0.05; ****p < 0.0001.

B. Random glucose level of suckling male Het mice on a mixed background at 6 weeks of age during an extended wean still reveals hyperglycemia, suggesting that weaning does not precipitate dysglycemia. **p < 0.01.

C-D. Weight gain as a proxy for overall health across genotypes, biologic sex, and age does not appear significantly altered between groups.

E. Insulin tolerance testing showing comparable peripheral insulin sensitivity in male and female WT and Het mice on a mixed background, similar to results in the previously published mixed background.

**Supplemental Figure 2. Senescent signatures are not evident in MafA^S64F/+^ female islets from C57 and mixed genetic backgrounds.**

qPCR showing senescence signature genes are not enriched in mixed (left) and C57 (right) background female Het islets at 6 weeks of age.

**Supplemental Figure 3. Binding ability of MAFA^WT^ and mutant MAFA^S64F^ are comparable.**

Gel shift assay of HeLa nuclear extract (NE) produced MAFA^WT^ and mutant MAFA^S64F^ bound to *INS* enhancer (known MAFA target), showing similar binding ability. WT comp, wild-type competitor. Mut comp, mutant competitor. Specificity of MAFA binding showed by super shift (SS) with MafA antibody.

**Supplemental Table 1**. Primer sequences and antibody sources

