## Supplementary figures and images for "Genetic background influences MAFA^S64F^-mediated diabetes penetrance in male mice"

### Supplemental Figures

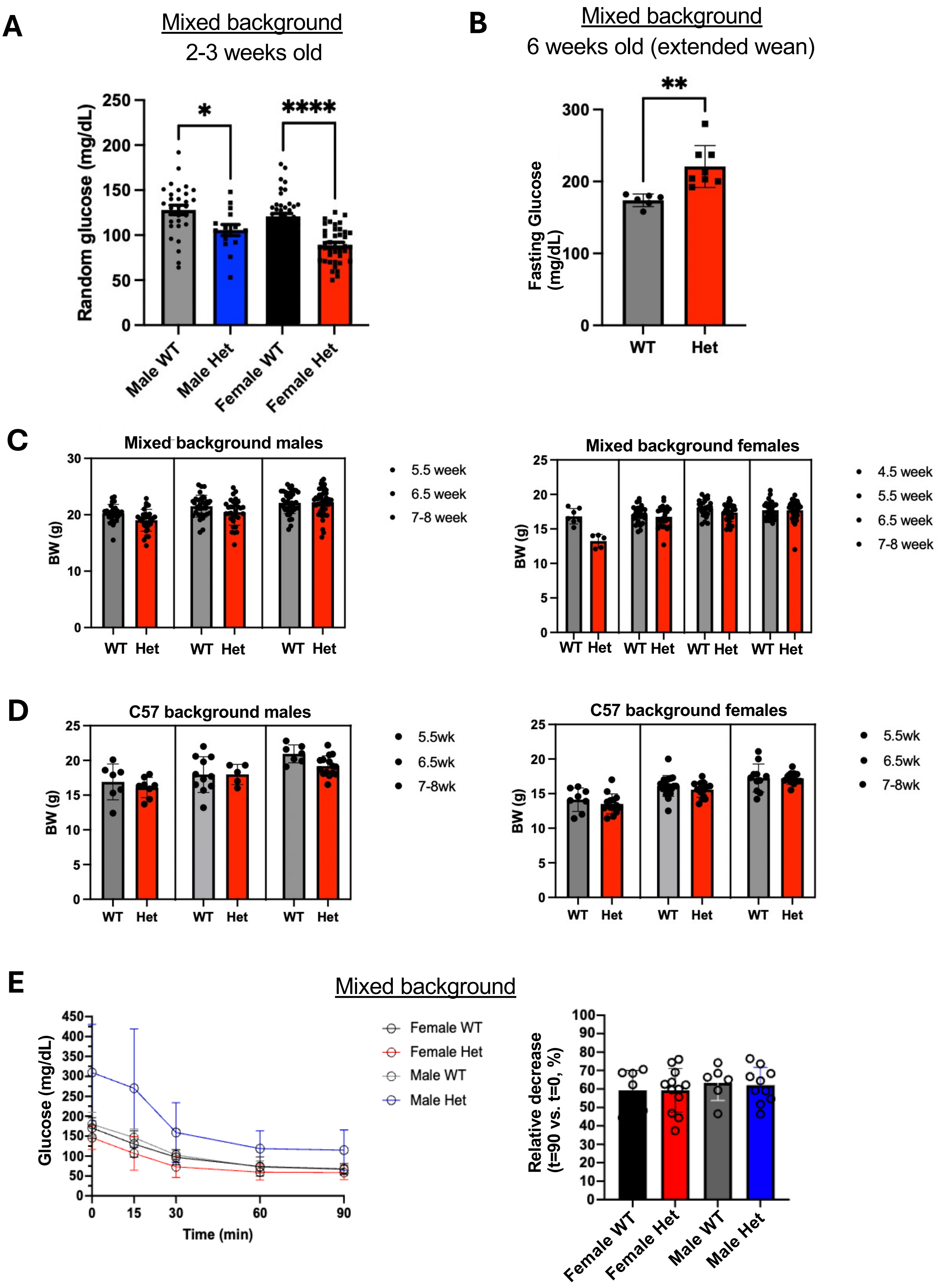


Supplemental figure 1

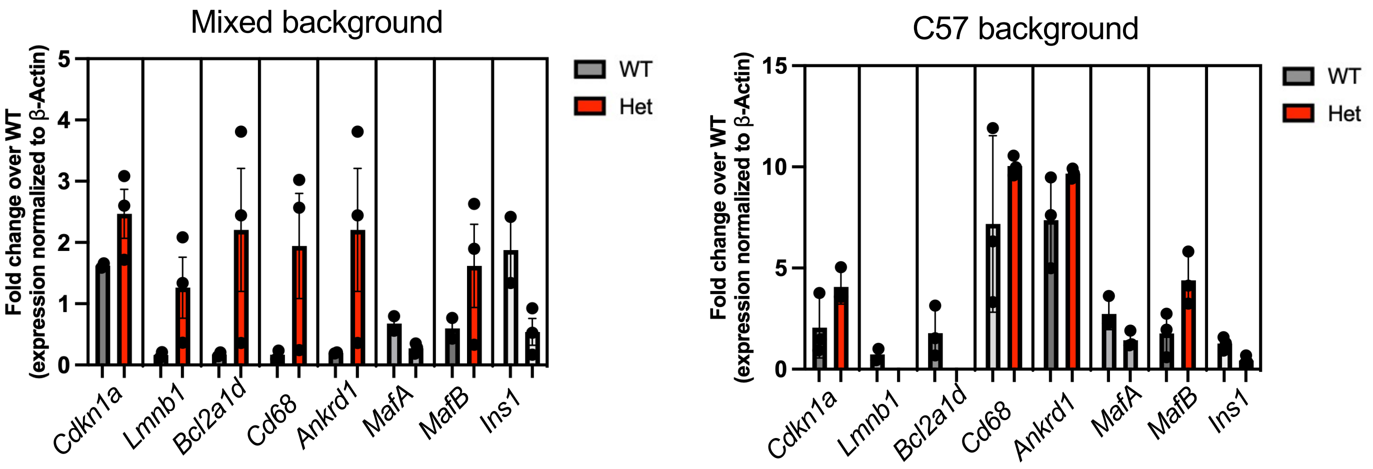


**Supplemental figure 2**

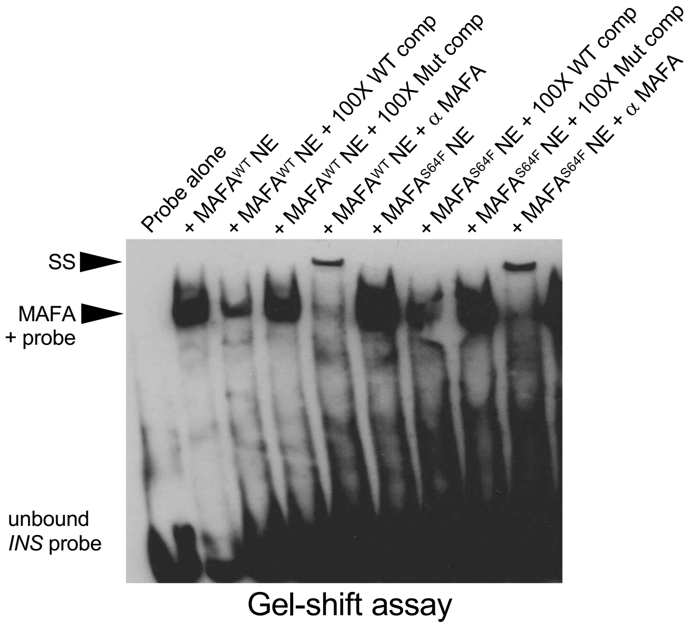


**Supplemental figure 3**
